# Local translation provides the asymmetric distribution of CaMKII required for associative memory formation

**DOI:** 10.1101/2022.03.28.486096

**Authors:** Nannan Chen, Yunpeng Zhang, Mohamed Adel, Elena A. Kuklin, Martha L. Reed, Jacob D. Mardovin, Baskar Bakthavachalu, K. VijayRaghavan, Mani Ramaswami, Leslie C. Griffith

## Abstract

How compartment-specific local proteomes are generated and maintained is inadequately understood, particularly in neurons, which display extreme asymmetries. Here we show that local enrichment of Ca^2+^/calmodulin-dependent protein kinase II (CaMKII) in axons of *Drosophila* mushroom body neurons is necessary for cellular plasticity and associative memory formation. Enrichment is achieved via enhanced axoplasmic translation of *CaMKII* mRNA, through a mechanism requiring the RNA-binding protein Mub and a 23-base Mub-recognition element in the *CaMKII* 3’UTR. Perturbation of either dramatically reduces axonal, but not somatic, CaMKII protein without altering the distribution or amount of mRNA *in vivo* and both are necessary and sufficient to enhance axonal translation of reporter mRNA. Together, these data identify elevated levels of translation of an evenly distributed mRNA as a novel strategy for generating subcellular biochemical asymmetries. They further demonstrate the importance of distributional asymmetry in the computational and biological functions of neurons.

## INTRODUCTION

CaMKII is crucial to behavioral plasticity across phyla [1-3]. The resting concentration of CaMKII is extraordinarily high, reaching 2% of total protein in the mammalian hippocampus [4], with most concentrated in synaptic regions. The 3’UTRs of both mammalian *CAMK2A* and *Drosophila CaMKII* contain regulatory information important for activity-dependent plasticity-related local translation in dendrites [5, 6] and presynaptic terminals respectively [7, 8]. While this acute modulation of CaMKII translation by neural activity has been described in multiple species, the mechanisms establishing, and indeed the function of, the impressive basal synaptic enrichment of CaMKII are completely unknown in any species.

## RESULTS

### CaMKII is enriched in axons via an active process

To visualize the extent of synaptic enrichment in *Drosophila melanogaster*, we expressed soluble GFP, which distributes in the cell via diffusion, under control of *CaMKII-GAL4* (Figure S1A), a driver transgene in the *CaMKII* locus, and compared its distribution to that of endogenous CaMKII. We focused on the mushroom body (MB), which is composed of Kenyon cells (KCs) with distinct somatic, dendritic and axonal compartments [9] (Figure 1A) whose plasticity is central to memory formation in *Drosophila* [10]. Figures 1B-C show the axon/soma ratio for soluble GFP is significantly lower than that of CaMKII suggesting that an active process regulates localization of CaMKII protein. This MB neuropil enrichment occurs specifically in KC axons, and is not due to CaMKII from extrinsic neuronal processes in the neuropil, as it is seen when the endogenous *CaMKII* gene is tagged with EGFP via CRISPR/Cas9 [11] exclusively in MBs (Figures 1D-F and S1B-C). Population of the synaptic region by trapping or stabilization of CaMKII can also be ruled out, since an EYFP::CaMKII fusion protein [12] expressed under control of GAL4 expresses at equivalent levels in cell bodies and axons (Figures 1G-H). These data suggest that non-coding regions (UTRs) of the *CaMKII* mRNA may control presynaptic accumulation of CaMKII protein.

**Figure 1.**
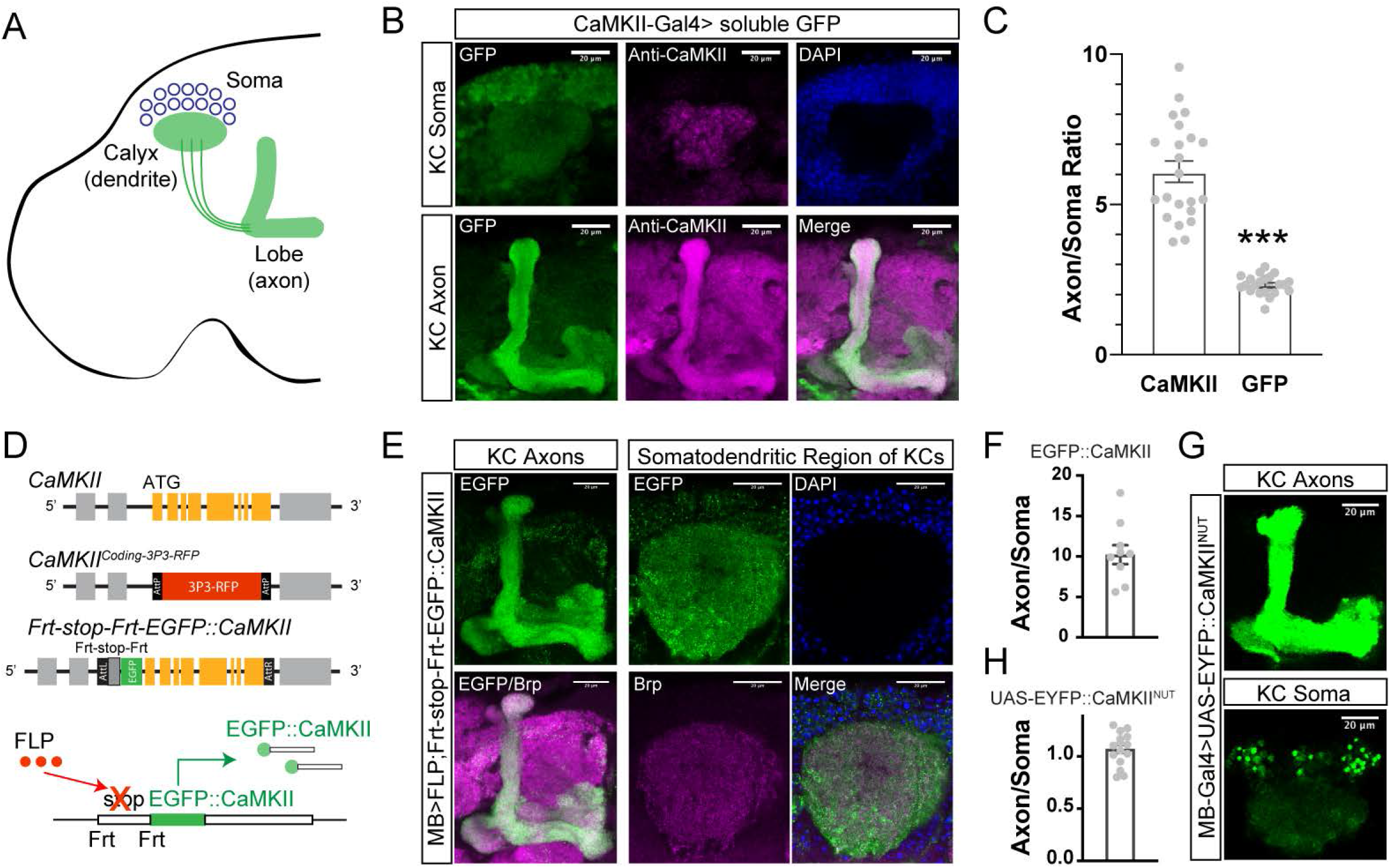
CaMKII is enriched in presynaptic regions relative to cell bodies by an active process. All transgenes were expressed with *VT030559-Gal4*, which is referred to in text/figures as *“MB-Gal4”* (A) Schematic diagram of mushroom body (MB) Kenyon cells. Dendritic processes, which receive inputs from the olfactory system, arborize adjacent to the somata, forming the “calyx”. Axonal processes extend ventrally to form the MB “lobes”. (B) Representative images of adult brain somatic (top) and MB axon (bottom) regions of animals expressing soluble GFP under control of *CaMKII-GAL4* visualized with GFP and anti-CaMKII. (C) Quantification of MB axon/soma ratio for CaMKII and soluble GFP. Data are mean ± SEM analyzed by Student’s t-test. Gray dots show individual values. N = 22 for each group. *** indicates p<0.001. (D) Schematic of *Frt-stop-Frt-EGFP-CaMKII* allele and recombination strategy. To construct a conditional EGFP::CaMKII fusion allele, we first replaced exons 1-8 of *CaMKII* with an attP flanked *3P3>RFP* cassette to make the *CaMKII*^*Coding-3P3-RFP*^ fly line. The RFP marker was then replaced by recombination of an attB-flanked *Frt-stop-Frt-EGFP* fused to the first eight exons of *CaMKII* to make the *Frt-stop-Frt-EGFP::CaMKII* fly strain. *Frt-stop-Frt* can be flipped out by cell-specific expression of Flp recombinase, allowing visualization of endogenous *CaMKII* protein levels. (E) Representative images of *MB>Flp* activation of EGFP::CaMKII protein expression in axons (left) and somatodendritic regions (right) stained with DAPI and anti-Brp. (F) Quantification of axon/soma ratio for EGFP::CaMKII. Data are mean ± SEM. Gray dots show individual values. N = 10. (G) Representative images of *MB-GAL4*-driven EYFP::CaMKII transgene lacking any *CaMKII* UTR. Axonal region (upper) and somatic region (lower) are shown. (H) Ratio of axon/soma EYFP was ca. 1, indicating no enrichment in axons. Gray dots show individual values. N = 16. Scale bars = 20 µm for each panel. See also Figure S1.

### The *CaMKII* 3’UTR contains multiple independent regulatory *cis*-elements

To ask if the *CaMKII* 5’ or 3’ UTR could mediate synaptic enrichment of protein in adult brain we generated animals with a GAL4-driven mMaple3 [13] coding sequence followed by either the long or short form of the 3’UTR (1990 and 123 bp respectively, produced by alternative polyadenylation [8]), or preceded by the 5’UTR of CaMKII (Figure 2A). All transgenes, including the control, which has no *CaMKII* UTR sequences, contain an SV40 polyadenylation site. Expression of these transgenes in KCs resulted in similar mMaple3 fluorescence in the somatic regions of all four lines (Figures S2A-B). Examination of the axonal processes of the MB, however, revealed a marked enrichment of mMaple3 protein in the MB lobes when the reporter was followed by the full 3’UTR (Figures 2B-C). Since mMaple3, like GFP, is a soluble cytosolic protein, somatically-synthesized protein can only reach axon terminals of the lobes by diffusion or transport. The ability of the long 3’UTR of CaMKII to concentrate mMaple3 at the synaptic terminal, without changing somatic protein levels, strongly suggests that the distal *CaMKII* 3’UTR is altering the location of protein synthesis.

**Figure 2.**
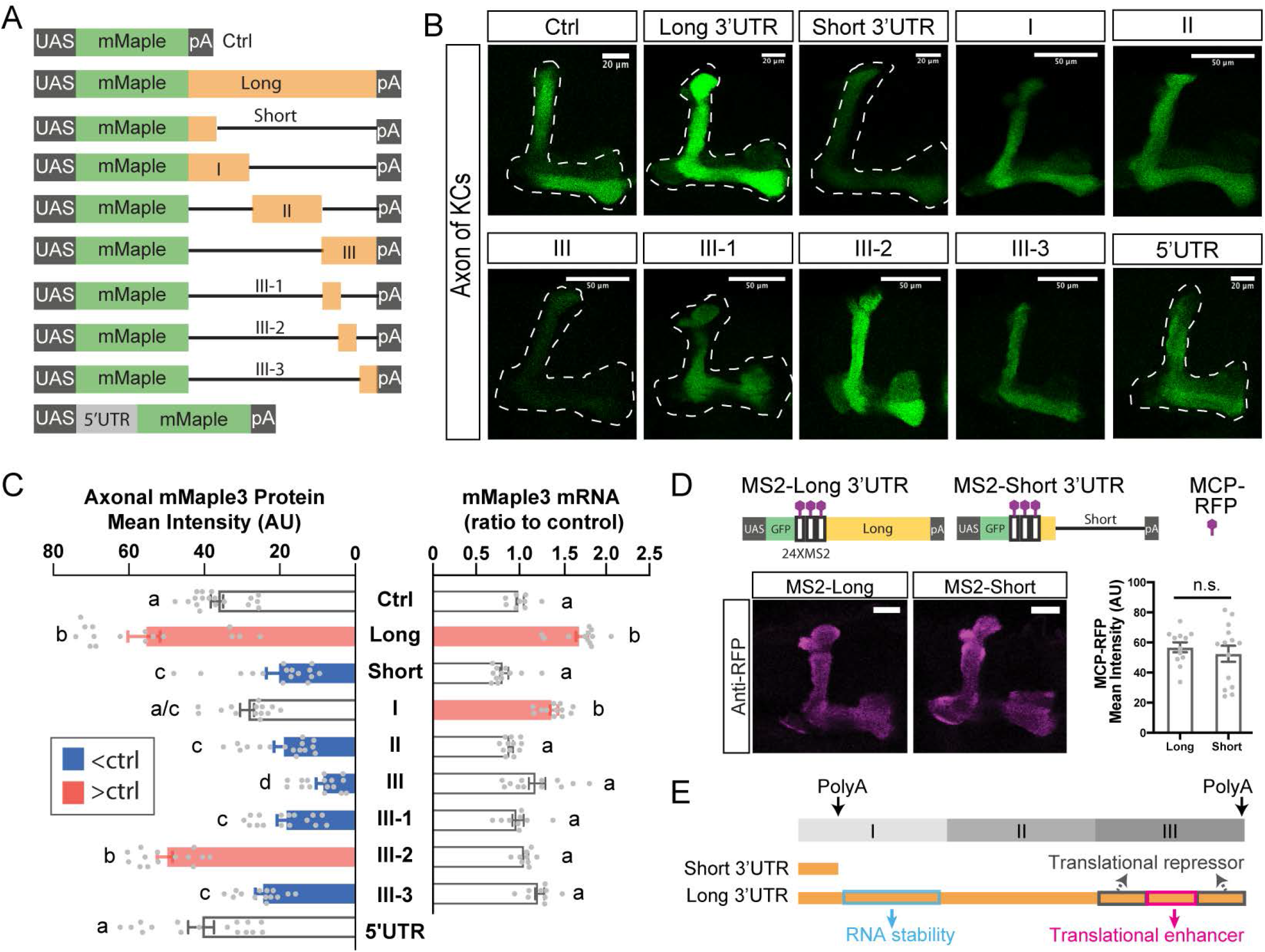
The distal *CaMKII* 3’UTR functions as the *cis*-element for axonal protein localization. (A) Reporter transgenes used to screen for *cis*-elements. All transgenes were expressed with *VT030559-Gal4*, which is referred to in text/figures as “MB-Gal4” (B) Representative images of mMaple3 protein in axons of *MB>mMaple3* animals. Scale bars = 20 µm. Dashed white lines indicate MB axons. (C) Left, mMaple3 protein levels in axons. N = 14-18. Right, qPCR of mRNA from adult *MB> mMaple3* heads using primers for *mMaple3*. N = 9-12. Red bars indicate significant increase relative to control, blue indicates decrease. (D) Schematic of protein/RNA reporters (top), and representative images of axonal MCP::RFP (bottom). Long and short UTR mRNAs localized to neuropil equivalently. N = 12-14. (E) Cartoon of *cis*-regulatory elements identified in this study. The long 3’UTR and fragment I both have higher levels of mRNA suggesting the presence of a stability element between the end of the short 3’UTR and the end of fragment I. The translational enhancer localizes to the III-2 region, while translational repressors localize to III-1 and III-3. Data are shown as mean ± SEM, analyzed by Student’s t-test or one-way ANOVA with Bonferroni post-hoc test as appropriate. Gray dots show individual values. Statistical differences are indicated by letters in panel C, with genotypes that are not significantly different having the same letter. n.s. indicates no significant difference in panel D. See also Figure S2.

Both the proximal and distal ends of the *CaMKII* 3’UTR are highly conserved in insects (http://genome.ucsc.edu), but only the function of the proximal and non-conserved middle regions have been previously studied due to incomplete annotation of the region leading to use of a truncated 3’UTR in previous studies [12]. The idea that 3’UTRs are modularized to execute multiple functions led us to explore the role of the conserved and unconserved pieces individually by making additional mMaple3 transgenes (Figure 2A) and examining their effects on protein and steady-state mRNA levels (Figures 2B-C). The proximal conserved region (fragment I) increased both mRNA and somatic mMaple3 compared to the SV40-only control (Figures 2C and S2A-C). However, these animals had about half the axonal mMaple3 of the long 3’UTR transgene (Figure 2C), indicating that, while the distal part of fragment I likely contains an mRNA stability element, (see also Figure 4D), it lacks a translation enhancer. Consistent with this, the non-conserved (II) and distal conserved (III) fragments had mRNA levels comparable to control and short 3’UTR transgenes, and lower protein expression in both soma and lobe compared to the full 3’UTR.

Interestingly, while the absolute levels of mMaple3 protein driven by fragment III were lower than the no-UTR control (suggesting the presence of translational repressor elements, see Figures 2C and S2B) the ratio of axonal to somatic protein appeared to be higher than for other transgenes, providing a hint that this region might harbor regulatory information for axonal translation. Additionally, this region is highly conserved across insect species, motivating us to subdivide it further. Division allowed us to map translational repressors to the proximal (III-1) and distal (III-3) parts of fragment III and revealed a strong axonal translation enhancer in fragment III-2 (Figures 2B-C). mRNA levels for all three were equivalent to control (Figures 2C and S2C). Both lobe (Figure 2C) and soma (Figure S2B) mMaple levels with the 286bp III-2 fragment were equal to those produced by the full length 3’UTR, suggesting that it contains the *cis*-element which is the major driver of axonal accumulation of CaMKII (Figure 2E).

### The distal *CaMKII* 3’UTR drives axonal protein enrichment via a translational mechanism

To determine if the enrichment of CaMKII protein was due to 3’UTR-dependent localization of mRNA or 3’UTR-dependent translation within the presynaptic compartment, we employed a second set of transgenes which contained, in addition to a myrGFP translation reporter, a 24-repeat MS2 string that allows detection of mRNA with transgenically-expressed MCP::RFP [14]. Figure 2D shows that there is no difference in the amount of axonal mRNA between transgenes containing short or full length 3’UTRs. This accumulation of MCP::RFP in axons was completely dependent on co-expression of an MS2 transgene (Figures S2E and S2G). As with the mMaple3 reporters, the presence of the distal 3’UTR significantly increased axonal myrGFP protein accumulation (Figures S2D and S2F), consistent with a role in translational regulation rather than mRNA localization. To ask if MB axonal processes are have the machinery for local translation, we expressed a GFP-tagged ribosomal subunit that has been shown to assemble with endogenous ribosomes [15] and found that it localizes to axons (Figure S2H), consistent with the presence of ribosomes. Taken together, these results support the idea that the distal part of the *CaMKII* 3’UTR specifies presynaptic accumulation of protein via a translational mechanism.

### RNA-binding protein Mub directs axonal protein accumulation

*Trans*-acting factors, including RNA-binding proteins (RBPs), function as the executors of the programs encoded in *cis*-elements [16]. To identify RBPs that might act on the *CaMKII* 3’UTR, we performed an *in silico* screen with RBPmap [17] and found that there were few RBPs predicted to bind only to the distal regions (Figures S3A-B). We assayed the function of these candidates by examining the effect of RNAis on reporter protein levels in animals expressing the *mMaple3-III-2* transgene. Only knock-down of *mushroom body expressed (mub)* [18], the widely-expressed single ortholog of the mammalian poly-C-binding-protein family, significantly decreased axonal mMaple3 protein expression (Figures 3A-B). Knockdown of RBPs that more generally bind the distal 3’UTR identified several candidate repressors (Figure S3C-D).

**Figure 3.**
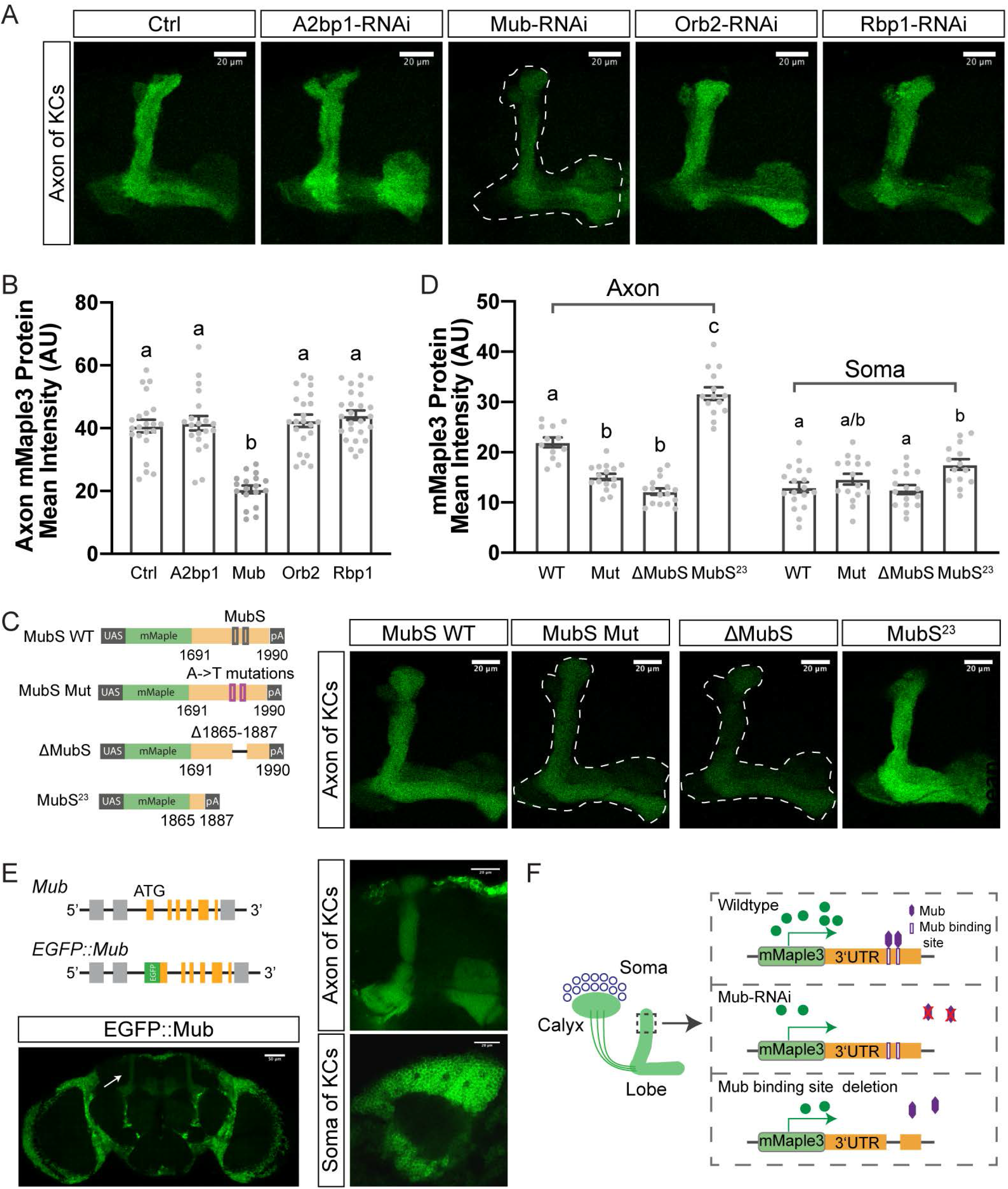
Mub is the *trans*-factor specifying CaMKII axonal enrichment. All transgenes were expressed with *VT030559-Gal4*, which is referred to in text/figures as “MB-Gal4”. (A) Representative single confocal sections of *MB>mMaple3-III-2* protein expression ± RNAi knock-down of RBPs predicted to bind to distal regions. (B) Quantification of MB axonal mMaple3 protein in *MB>mMaple3-III-2* ± RNAi brains. N = 18-26. (C) Left, schematic of transgenes with Mub site manipulations. Right, representative pictures of mMaple3 in *MB>mMaple3-MubS* axons. (D) Quantification of axonal and somatic mMaple3 protein levels from Mub site mutant transgenes. Loss of Mub-binding reduced mMaple3 in axons, while leaving somatic levels unchanged. The isolated 23bp Mub binding site elevated axonal protein over the level of control, and slightly elevated somatic levels, consistent the presence of repressor elements in the control UTR fragment. N = 12-18.1 (E) Diagram of CRISPR-engineered *EGFP::Mub* allele. Images show EGFP for whole brain, MB axons and soma. (F) Cartoon summary of Mub manipulations. Red X indicates no expression of Mub. For all panels: Scale bars = 20 µm. Dashed white lines indicate the MB axons. Data are mean ± SEM and quantified by one-way ANOVA with Bonferroni post-hoc test. Gray dots show individual values. Statistical differences are indicated by letters; groups that are not significantly different have the same letter. See also Figure S3.

While the RNAi results were consistent with Mub directly regulating reporter accumulation via the *CaMKII* 3’UTR, it was also possible that the decrease in mMaple3 protein was due to a more general or indirect function of Mub. To determine if the effect of Mub knockdown was direct, we mutated predicted Mub-binding sites within the 3’UTR and asked if that affected mMaple3 protein levels in axons. We used a transgene that contained both fragments III-2 and -3 since Mub has two binding sites in fragment III, one of which spans the III-2/3 junction. Both A->T point mutations in the two sites, or deletion of the 23bp containing the sites, decreased mMaple protein expression in axons (Figures 3C-D). Notably, the 23bp element alone was sufficient to raise expression over the level of the WT fragment (Figures 3C-D). Importantly, somatic mMaple3 levels were not significantly changed by loss of Mub sites (Figure 3D). We speculate that this, as well as the high activity of the isolated 23bp fragment, is due to the presence of repressor sequences in region III-3 that normally act to block translational stimulation by Mub in the cell body.

While early studies [18] and our *Mub-GAL4* line (Figure S1D) suggested that the *mub* gene is transcribed in MB, it was important to determine where the Mub protein was localized within the MB. We used CRISPR/Cas9 [11] to fuse EGFP in-frame to the N-terminal of the Mub coding region. Consistent with *Mub-GAL4*’s expression pattern, EGFP::Mub protein was widely-expressed, and almost exclusively somatic in the adult brain. The one remarkable exception was the MB, where EGFP::Mub can be seen in axonal processes (Figure 3E). These data suggest that Mub has a unique role in CaMKII axonal translation in KCs (Figure 3F).

### Mub sites in the *CaMKII* 3’UTR mediate formation of the basal proteome and support associative memory formation

To explore the function of the distal 3’UTR in the context of the endogenous *CaMKII* locus, we used CRISPR/Cas9 [11] to replace the entire 3’UTR with an attP-flanked 3XP3-RFP recombination cassette, creating *CaMKII*^*UDel*^ (Figure 4A). We then replaced 3XP3-RFP using attB-flanked 3’UTR sequences to engineer full, short and Mub-site mutant 3’UTR lines (*CaMKII*^*ULong*^, *CaMKII*^*UShort*^ and *CaMKII*^Δ*MubS*^) (Figure 4A). Both *CaMKII*^*UShort*^ and *CaMKII*^Δ*MubS*^ had drastically decreased axonal (Figures 4B-C) and total head CaMKII protein (Figures S4A-B) compared to *CaMKII*^*ULong*^. Levels of mRNA for *CaMKII*^*ULong*^ and *CaMKII*^Δ*MubS*^ were not different (Figure 4D). *CaMKII*^*UShort*^ mRNA was ∼10% lower, likely due to loss of a stability element (see Figure 2E), but this decrease is not enough to account for the ∼75% decrease in protein. These data suggest that loss of the 23bp Mub site renders animals incapable of generating high basal axonal CaMKII protein despite the fact that total mRNA levels are normal (Figure 4D).

**Figure 4.**
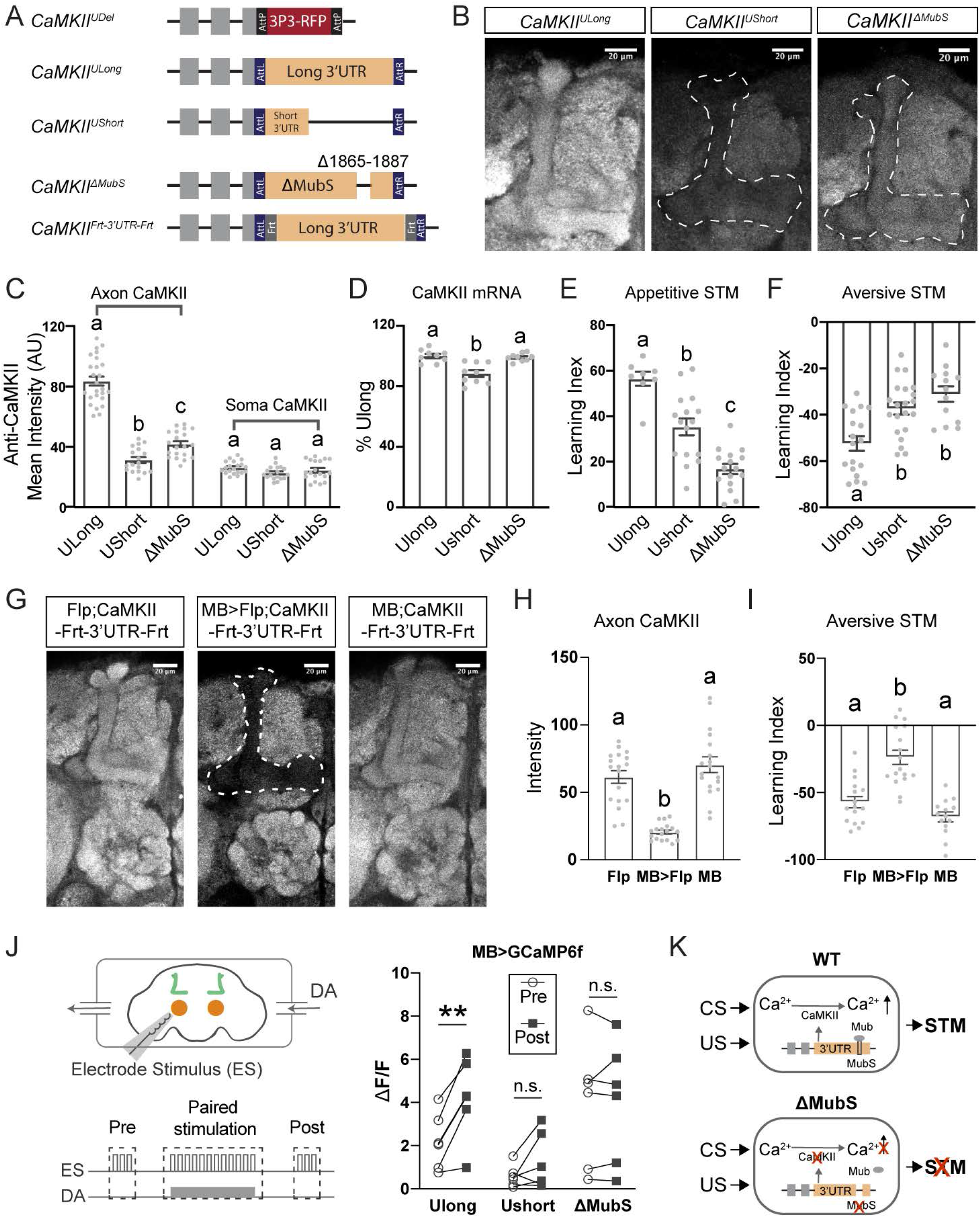
Loss of Mub-binding sites in the *CaMKII* 3’UTR impairs plasticity and memory formation. All transgenes were expressed with *VT030559-Gal4*, which is referred to in text/figures as “MB-Gal4” (A) Diagram of CRISPR-engineered *CaMKII* alleles. *CaMKII*^*Ude*l^ has the entire 3’UTR replaced by a recombination site-flanked 3P3>RFP transgene; other lines were generated from this by recombination. Endogenous *CaMKII* exons are indicated in gray, and replaced regions in color. (B) Representative images of MB axon CaMKII protein levels in *CaMKII*^*ULong*^, *CaMKII*^*UShort*^and *CaMKII*^Δ*MubS*^. Dotted white lines indicate axonal region. Scale bars = 20 µm. (C) Anti-CaMKII staining in MB axons is reduced in *CaMKII*^*UShort*^ and *CaMKII*^Δ*MubS*^. N = 18-25. (D) *CaMKII* mRNA levels measured by qPCR of total head mRNA are slight lower in *CaMKII*^*UShort*^, but not affected by loss of Mub sites. N = 9 for each group. (E) Appetitive short-term memory is disrupted in *CaMKII*^*UShort*^ and *CaMKII*^Δ*MubS*^. N = 8-16. (F) Aversive short-term memory is disrupted in *CaMKII*^*UShort*^ and *CaMKII*^Δ*MubS*^. N = 13-21. (G) Representative pictures of MB-specific deletion of the 3’UTR. Dotted white line indicates MB axons. Scale bars = 20 µm. (H) CaMKII protein is reduced exclusively in MB axons when the 3’UTR is specifically knocked out in MB. N = 18 per group. (I) Aversive short-term memory is disrupted when the 3’UTR is deleted specifically in MB. N = 14-17. (J) Left, diagram of PDP paradigm. Antennae lobe stimulation trains (ES, white bars) and dopamine perfusion (DA, solid gray bar) are paired for 1 min. Right, Post-pairing calcium response increased significantly in *CaMKII*^*ULong*^, but not in *CaMKII*^*UShort*^ or *CaMKII*^Δ*MubS*^. Circles and squares show individual values. Data were analyzed by paired-T-test, n.s. indicates no significant difference, ** indicates p<0.01. (K) Model. Potentiation of calcium responses by association requires high levels of synaptic CaMKII. When Mub sites are lost, presynaptic CaMKII protein is reduced, and MB plasticity and memory formation are blocked. Except for panel J, data are mean ± SEM analyzed by one-way ANOVA with Bonferroni post-hoc test. Gray dots show individual data points. Statistical differences are indicated by letters; groups that are not significantly different have the same letter. See also Figure S4.

We next asked if CaMKII accumulation in axons was needed for MB plasticity by assaying short-term associative memory in 3’UTR-mutant lines. Loss of the distal 3’UTR significantly impaired both appetitive (Figure 4E) and aversive (Figure 4F) associative memory formation. This defect mapped to the Mub-binding element since animals lacking only the 23bp site were almost totally unable to learn (Figures 4E-F). To determine if this plasticity defect was a result of loss of axonal CaMKII specifically in KCs, we utilized a conditional allele, *CaMKII*^*FRT-3’UTR-FRT*^ (Figure 4A) and removed the 3’UTR of the gene exclusively in KCs by expression of FLP recombinase. These animals, which have normal synaptic CaMKII in all cells except for KCs (Figures 4G-H and S4G), showed significant decreases in aversive associative memory formation (Figure 4I). Shock and odor detection were similar in these flies (Figures S4C-F and S4H-I).

A critical early event of associative memory formation in the α’ lobe of MB is potentiation of odor responses, postulated to be a memory trace or engram [19]. To determine if the basal CaMKII proteome was required for this cellular process, we utilized an *ex vivo* assay for pairing-dependent plasticity (PDP), a paradigm similar to long-term enhancement [20, 21] in which stimulation of the antennal lobe olfactory inputs (the conditioned stimulus) is temporally paired with dopamine application (the unconditioned stimulus). Potentiation of the response to antennal lobe stimulation in the α’3 compartment was defective in both *CaMKII*^*UShort*^ and *CaMKII*^Δ*MubS*^ (Figure 4J). Importantly, the initial response of KCs to acetylcholine, the transmitter used by olfactory inputs to the MB, was not altered by UTR mutations (Figure S4J-S4M). These results demonstrate that the enrichment of CaMKII in the presynaptic proteome by a Mub-dependent mechanism is required for the plastic changes in odor responses that are a signature of associative memory formation (Figure 4K). This underscores the critical role of high levels of axonal CaMKII in the computational processes carried out in this compartment.

## DISCUSSION

Local protein synthesis at synapses has been studied extensively in the context of specialized processes like activity-dependent plasticity and axon guidance [22]. Recent theory and experimental work [23, 24], however, suggests that local translation occurs much more generally and may be used to establish differential proteomes in functionally-specialized subcellular regions. In this study we resolve two long-standing questions about CaMKII: how and why it achieves extraordinary levels in axons. We demonstrate that resting adult levels of CaMKII protein are translationally accrued, and that the high levels in this compartment form a computational scaffold critical for formation of associative memory and the cellular memory trace. While previous studies using mutants and RNAi have shown a role for CaMKII in plasticity, our manipulations of the 3’UTR, which do not affect somatic kinase levels, establish the necessity of synaptic enrichment. This enrichment requires *cis-*elements present only in the long form of the 3’UTR and Mub, the *Drosophila* poly-C-binding-protein homolog demonstrating a new, activity-independent function for the *CaMKII* 3’UTR.

Activity-dependent translation [5, 7, 12, 25, 26] and differential polyadenylation [24, 26] are ancient conserved features of CaMKII mRNAs. For mammalian *CAMK2A*, early work in which the 3’UTR was deleted demonstrated its requirement for mRNA stability and dendritic localization, and also for protein accumulation and activity-dependent synthesis [6]. A handful of studies attempted to identify *cis*-elements regulating dendritic *CAMK2A* mRNA localization and transport [27-29], but there is as yet no information on 3’UTR *cis-*elements controlling translation, though *in silico* prediction suggests that the *CAMK2A* 3’UTR may have polyC-binding protein motifs.

At the *Drosophila* larval neuromuscular junction, we and others have shown that the *CaMKII* 3’UTR controls activity-dependent synthesis of CaMKII [7, 12, 26, 30]. The fact that the rodent *CAMK2A* 3’UTR can support activity-dependent protein synthesis in the fly [12] suggests that there will be shared mechanisms for this aspect of CaMKII regulation. But while there are many similarities between mammals and flies, there are also differences. In *Drosophila*, the 3’UTR appears to have little effect on mRNA localization, and only a small effect on stability that is ascribable to a proximal *cis-*element. How *CaMKII* mRNA reaches synapses in *Drosophila* is yet to be determined, but the differences in localization mechanism may reflect the ca. 100-fold difference in distances that mRNAs need to travel to reach synapses.

The ability of Mub, which is present at low levels in MB axons and at high levels in MB and other cell bodies, to specifically regulate axonal accumulation of CaMKII protein without affecting somatic protein levels suggests several models. One possibility is that MB axons have either compartment-specific translational machinery or a distinct set of auxiliary proteins that allow Mub to regulate axonal ribosomes [31, 32]. The presence of Mub protein in MB axons, but not in other neuropils, may indicate the existence of unique translational complexes in that compartment. Another possibility is that Mub is a general translation enhancer, but MB soma contain repressor proteins that locally inhibit its actions. This would be consistent with our finding that there are *cis* elements that appear to act as general repressors in the *CaMKII* 3’UTR. While these ideas remain speculative, the robust interaction of Mub with *CaMKII* provides an opportunity to deepen our understanding of how local protein synthesis can shape neuronal function and build the synaptic proteome.

## ACKOWLEDGEMENTS

We thank Ed Dougherty in the Brandeis Imaging Facility for assistance. Stocks obtained from the Bloomington Drosophila Stock Center (NIH P40 OD018537) were used in this study. This work was supported by NIH R37 NS112810 (to LCG), NIH R01 DA043195 (to LCG), a Science Foundation Ireland Investigator Programme Grant and Wellcome Trust-HRB-SFI Grants (to MR).

## AUTHOR CONTRIBUTIONS

LCG and NC conceived the study and designed the experiments. NC, YZ and BB constructed fly lines. NC did all immunohistochemical analysis and qPCR. EAK did immunoblotting and analysis. NC, JDM and MLR did behavioral experiments. MA did electrophysiological experiments. All authors assisted with data analysis and interpretation. NC and LCG wrote the manuscript with input from all the authors.

## DECLARATION OF INTERESTS

The authors declare no competing interests.

## METHODS DETAILS

### Fly strains

Flies (*Drosophila melanogaster)* were raised on dextrose/cornmeal/yeast food at 25°C with a 12h:12h light-dark cycle. Male and female flies were collected at eclosion and aged for 2-4 days before performing immunohistochemical experiments and for 7-10 days for behavior. For GCaMP experiments, only female flies (4-8 days old) were used due to the size and clarity of their brains. *UAS-mCD8::GFP* flies [33] were a gift from Dr. Liqin Luo (Stanford University), and *EYFP::CaMKII*^*NUT*^ flies [12] from Dr. Sam Kunes. *VT030559-GAL4, A2bp1-RNAi* (110518), *Mub-RNAi* (105495), *Orb2-RNAi* (107153) and *Rbp1-RNAi* (110008) flies were obtained from VDRC Stock Center. *UAS-GFP::RPL10, UAS-MCP::RFP, UAS-GFP, 20XUAS-GCaMP6f, UAS-Flp, Mub*^*[MI08161]*^*/TM3SbSer* (43942) and *CaMKII* ^*[MI03976]*^ (60770) flies were ordered from Bloomington *Drosophila* Stock Center. All transgenic, MiMIC conversion and CRISPR injections were performed by Rainbow Transgenics (Camarillo, CA).

### Creation of mMaple3-UTR and myrGFP-MS2-UTR transgenic lines

For the mMaple3 lines, all 3’UTR fragments were amplified from the *Canton-S* wild type fly genome. The list of PCR primers is in Table S1. The mMaple3 plasmids were a gift of Dr. Margret Stratton (UMASS Amherst). To make the *UAS-mMaple3-long 3’UTR* line, the mMaple3 fragment and the long 3’UTR fragment were amplified and then inserted into the pUAST-attB plasmid (Addgene, 8489bp) using the Gibson assembly method. The mMaple3 fragment was followed by the long 3’UTR fragment. For other 3’UTR lines, we used the same mMaple3 fragment and pUAST-attB plasmid. For the *UAS-mMaple3* control line, only the mMaple3 sequence was put into the pUAST-attB plasmid. For the *UAS-5’UTR-mMaple3* fly strain, the zygotic 5’UTR was amplified by PCR and inserted before the mMaple3 fragment. All plasmids were checked by sequencing. Plasmids were injected into *phiC31-attP* flies (Bloomington Stock Center #79604) which have attP sites on the second chromosome to allow targeted integration. The progeny of injected flies was screened by *w*^*+*^ red eye marker, and then checked by PCR and sequencing.

Transgenic *myrGFP-MS2-UTR* lines were created by site-specific insertion of pJFRC-myrGFP-MS2-long-3’UTR and pJFRC-myrGFP-MS2-short-3’UTR plasmids into *phiC31-attP* docking site (Bloomington Stock Center #79604). To construct these plasmids, long or short 3’UTR sequences were amplified from *Canton-S* flies and inserted into the XbaI digested pJFRC12-10XUAS-IVS-myrGFP plasmid (Addgene #26222) using the Gibson assembly method. The MS2 sequence was amplified from the TRICK plasmid obtained from Dr. Jeffrey A. Chao (Friedrich Miescher Institute for Biomedical Research, Basel, Switzerland) and cloned between the GFP and the *CaMKII* UTR sequence to create pJFRC-myrGFP-MS2-long-3’UTR and pJFRC-myrGFP-MS2-short-3’UTR plasmids. The PCR primers are listed in Table S1. The injections of these plasmids were performed by the fly facility at Bangalore Life Science Cluster (BLiSC).

### Creation of endogenous *CaMKII* UTR deletion fly and *CaMKII*^*coding-3P3-RFP*^ fly

To make the endogenous *CaMKII* UTR deletion fly (*CaMKII*^*Udel*^), we designed a guide RNA which recognize a site in the *CaMKII* 3’UTR region (Table S2). The guide RNA was cloned into pU6 plasmids (Addgene, #45946). To replace the full 3’UTR region precisely, we made a donor plasmid which contained two homology arms: the left arm was a 2 kb length fragment which preceded the stop codon TAA, and the right arm is a 2 kb-length fragment which followed the last nucleotide of the full 3’UTR. Between the two arms was an inverted-attP flanked 3xP3-RFP fragment (Addgene, #80898), which was used as a marker to check insertion (CaMKII Udel-homologous arms plasmid in supplemental plasmids). A mixture of guide RNA plasmid and the donor plasmid was injected into Cas9 flies (*y,sc,v; nos-Cas9/CyO; +/+*). By the same strategy, we designed two guide RNAs (Table S2) which recognize the beginning and Exon 8 of CaMKII coding region separately, and a donor plasmid (CaMKII coding-homologous arms plasmid in supplemental plasmids) to make the *CaMKII*^*coding-3P3-RFP*^ fly.

After crossing to *ci*^*D*^*/ey*^*D*^ flies, the F1 progeny with the RFP marker were selected as candidates. Correct integrations were confirmed by PCR and sequencing with primers which bind to outside the regions of the integrated junctions.

### Creation of endogenous *CaMKII* UTR knock-in fly lines

To make *CaMKII*^*ULong*^ flies, the long 3’UTR sequence was amplified and flanked by two inverted-attB sites, then cloned into the pBS-KS-attB2 plasmid (Addgene, #62897). This plasmid was injected into *CaMKII*^*Udel*^ flies, with plasmids that expressed phiC31 recombinase. F1 progeny without RFP marker were selected as candidates, and further confirmation by PCR and sequencing were performed. Using the same strategy, we amplified the short 3’UTR sequence and made *CaMKII*^*UShort*^ flies. For the *CaMKII*^*ΔMubS*^ fly, we deleted 23 bp containing the two Mub binding sites from the long 3’UTR sequence. For the *CaMKII*^*Frt-3’UTR-Frt*^ fly, the whole 3’UTR flanked by two Frt sites were cloned. The recombination plasmids are listed in the supplemental plasmids. All flies have been checked by PCR and sequencing with primers outside of the insertion fragment.

### Creation of *Frt-stop-Frt-EGFP::CaMKII* fly line

For the *Frt-stop-Frt-EGFP::CaMKII* fly line, we amplified the stop sequence which is flanked by two Frt sites, *EGFP* sequence, and *CaMKII* sequence between two attp sites of *CaMKII*^*coding-3P3-RFP*^ fly. These fragments were assembled in order and cloned into the pBS-KS-attB2 plasmid (Frt-stop-Frt-EGFP-CaMKII plasmid in supplemental plasmids). This plasmid was injected into *CaMKII*^*coding-3P3-RFP*^ flies, with plasmids that expressed phiC31 recombinase. F1 progeny without RFP marker were selected as candidates, and further confirmation by PCR and sequencing were performed.

### Creation of endogenous *CaMKII-GAL4, Mub-GAL4* and *EGFP::Mub* flies

To make the endogenous *CaMKII-GAL4* fly strain, the phase 1 T2A-GAL4 plasmids (Addgene, #62897) and pBS130 plasmids (Addgene, #26290) which encode phiC31 integrase were injected into *CaMKII*^*[MI03976]*^ flies. Progeny were crossed to *yw, UAS-mCD8::EGFP* flies to check for GAL4 insertion. Male flies with yellow marker were selected as candidates and checked for GFP expression to obtain the insertion lines which were in the correct orientation. For *Mub-GAL4* flies, we used the same strategy, and phase 0 T2A-GAL4 plasmids (Addgene, #62896) were injected into the *Mub*^*[MI08161]*^*/TM3, SbSer* flies.

For *EGFP::Mub* flies, we designed a guide RNA which recognized the beginning of *Mub* (Table S2) and a donor plasmid (EGFP::Mub plasmid in supplemental plasmids). The guide RNA was cloned into pU6 plasmids (Addgene, #45946) and injected into Cas9 flies with the donor plasmid. Correct integrations were confirmed by PCR and sequencing with primers which bind outside the regions of the integrated junction.

### Dissection and immunohistochemistry

Fly brains (males and females, 2-4 days old) were dissected in cold Schneider’s Insect Medium (Sigma, S0146). To minimize any systemic error caused by the order of dissection and length of freezing, all fly lines were frozen on ice at the same time. Then we dissected one fly from each line, and performed the second round dissection. After dissection the brain samples were fixed in 4% PFA solution for 45 mins at room temperature, then washed 3×30 mins in 0.5% Triton-PBS (PBST) solution. For the *mMaple3-UTR* lines, the samples were mounted anterior side up in Vectashield (Vector Labs, Burlingame, CA) mounting medium after washing. For brains that needed antibody staining, after fixation and washing, the samples were blocked in 10% normal goat serum solution for 1 hour, and incubated in primary antibody solutions for 2-3 days. Primary antibody solutions were removed, and the samples were washed in PBST solution for 3×30 mins. After that the samples were incubated in secondary antibody solutions overnight. After 3×30 mins PBST washing, the same mounting protocol was performed afterwards.

For CaMKII staining the fixation time was shortened to 30 minutes. The primary antibodies used were: mouse anti-CaMKII (1:10,000, Cosmo CAC-TNL-001-CAM), rabbit anti-DsRed (1:150, Takara, 632496) and rabbit anti-GFP (1:1000, Thermo Fisher A-11122). DAPI was used at 1:1,000 (Invitrogen, D1306). Alexa Fluor 488 anti-mouse antibody (Invitrogen, A28175) and Alexa Fluor 635 anti-rabbit antibody (Invitrogen, A-31576) were used as secondary antibodies at 1:200 dilutions.

### Image processing and intensity analysis

Images were taken using Leica SP5 confocal microscope under a 20x objective lens, and then analyzed using ImageJ Fuji software [34]. For comparisons between several lines, all images were taken at the same laser strengths, gains and all other settings. To quantify the intensity of axonal MB regions, we selected all consecutive slices which contain MB lobes, chose the middle slice as the representative picture and outlined all lobes as the region of interest (ROI). We quantified the mean intensity of that region to be analyzed further. To quantify the intensity of somatic regions, we checked all consecutive slices that contain the cell bodies, chose the middle slice, outlined all the cell bodies in the slice and quantified mean intensity of the outlined ROI. Further analysis of the intensity was performed using GraphPad (GraphPad Software, San Diego, CA) program.

### Real-time PCR experiments and quantification

For each genotype, RNA was extracted from approximately 100 males and female fly heads using the Trizol Reagent (Thermo Fisher). The age of the flies was 2-4 days. RNA samples (1 ug) from each genotype were reverse-transcribed using PrimeScript 1st strand cDNA Synthesis Kit (Takara). Quantitative real-time PCR experiments were performed using TB Green Advantage qPCR premixes kit (Takara) on the Thermocycler (Eppendorf Realplex2). The primers for CaMKII, mMaple3 and ribosomal gene rp49 (Table S3) were ordered from Eton company. Quantification of rp49 was used for normalization.

### Western Blotting and quantification

Flies heads (10 flies per sample, males and females) were ground in loading buffer (NuPAGE™ LDS Sample Buffer), heated at 70°C for 10 minutes, and then 2-Mercaptoethanol was added to the samples. Proteins were separated by SDS-PAGE (NuPAGE™ Bis-Tris Protein Gels, Invitrogen), and transferred to nitrocellulose membrane (Amersham) in transfer buffer (NuPAGE). The membranes were blocked for 1 hour (blocking buffer for fluorescent western blotting, Rockland Immunochemicals), and then incubated with mouse anti-CaMKII (1:400, Cosmo) and mouse anti-Actin (1:1000, Millipore) solution overnight. Secondary antibody was anti-mouse IgG DyLight™ 680 conjugated pre-adsorbed antibody (1:5000, Rockland Immunochemicals).

Membranes were scanned on a ChemiDoc™ Touch Imaging System with Image Lab™ Touch Software (Bio-Rad), and quantified with Image Lab™ Software from Bio-Rad. Intensity of bands was measured with the Volume Tool and calculated using the background-adjusted volume. Intensity of CaMKII band was normalized to the actin signal in the same lane.

### Appetitive and aversive associative learning assays

Appetitive and aversive associative olfactory learning experiments were performed in an environment room in red light at 25°C and 70% humidity. Age of the flies (males and females) was between 7-10 days. The two odors used were 4-methylcyclohexanol (MCH) (Sigma, 153095) and 3-octanol (OCT) (Sigma, 218405) which were diluted in mineral oil at 10% concentration.

For the appetitive learning, filter papers were prepared with 2M sucrose, with blank filter papers used as control. Before training, the flies were starved to 10% mortality. 50-100 flies were loaded into a vial and given approximately 10 min acclimation in the apparatus. After acclimation, they were exposed to one odor (CS+) with the sucrose paper for 1 minute, and then to the other odor (CS-) with the blank paper for 1 minute with a 1-minute interval. For aversive leaning, the CS+ was exposed with 60 V electric shock for 1 minute, and then CS-for 1 minute with 1-minute interval.

After training, the flies were allowed to choose between the two odors for 2 minutes. A preference index (PI) was calculated as [(number of flies in CS+)-(number of flies in CS-)]/ [(number of flies in CS+) + (number of flies in CS+)]. For each genotype, two groups of flies were trained, one with MCH as CS+ and the other with OCT as CS+. The final learning index was calculated as the average of PIs from the two reciprocal trials.

### Functional calcium imaging

All imaging experiments were performed using a dissected brain preparation. Briefly, female, 4-8 day old brains were dissected in cool HL3.1 and placed into an imaging chamber. The dissected brains were allowed to recover for about 10 mins before stimulation. An electrode was used to stimulate olfactory projection neurons by putting the end to the antennal lobe. Perfusion flow was established over the brain with a gravity-fed ValveLink perfusion system (Automate Scientific, Berkeley, CA). Imaging was performed using an Olympus BX51WI fluorescence microscope (Olympus, Center Valley, PA) under 40x water immersion lens. The calcium signals were recorded using a charge-coupled device camera (Hamamatsu ORCA C472-80-12AG).

Hemolymph-like saline (HL3.1) consisting of (in mM) 70 NaCl, 5 KCl, 0.1 CaCl2, 20 MgCl2, 10 NaHCO3, 5 trehalose, 115 sucrose, 5 HEPES; pH 7.1-7.2 [35] was continuously perfused into the imaging chamber. Dopamine and acetylcholine were purchased from Sigma– Aldrich (St Louis, MO). For the *ex vivo* learning paradigm, the brains were first given 5 stimulation trains (20 pulses at 100 Hz; pulse width is 1 millisecond, and interpulse interval is 9 milliseconds) with 15 seconds intertrain interval to establish baseline. Calcium responses to the 5 trains were averaged to calculate the “Pre” responses. The brain was allowed to rest for 45 seconds before induction. Pairing-dependent plasticity (PDP) [21] was induced by pairing dopamine (10 micromolar) perfusion with 12 trains of AL stimulation (5 seconds inter-train interval; the same train profile as described above). After 15 minutes we repeated the 5 trains of antennal lobe stimulations to measure the “Post” response. For acetylcholine perfusion experiments, the dissected brains were exposed to 4 mM or 10 mM acetylcholine for 1 min, and then washed with HL3.1 saline.

The alpha prime tips (α’) of the MBs were selected as regions of interest. Signals were analyzed using custom software developed in ImageJ (National Institute of Health, Bethesda, MD). The percent change of fluorescence was calculated as ΔF/F = (Fn – F_0_)/F_0_ × 100%, where Fn is the fluorescence at time point n, and F_0_ is the fluorescence at time 0. The averages of maximum percent change were determined as the response value for each trial.

Statistical analyses were performed using Matlab (The MathWorks). For PDP, a paired t-test was used to determine statistical significance between pre and post responses. For acetylcholine response, one-way ANOVA followed by Bonferroni post-hoc test was used to determine the significance among groups.

### Statistics

We used GraphPad (GraphPad Software, San Diego, CA) to make histograms and analyze data except as noted. For comparison among three or more groups, we performed one-way ANOVA, and followed with Bonferroni post-hoc test to compare difference between groups. For comparison between two groups, we used the Student T-test. P<0.05 is indicated as *, p<0.01 is indicated as **, and p<0.001 is indicated as ***.

### Data availability

The data generated and analyzed during this study are available from the corresponding author on request.

---

**Figure S1.**
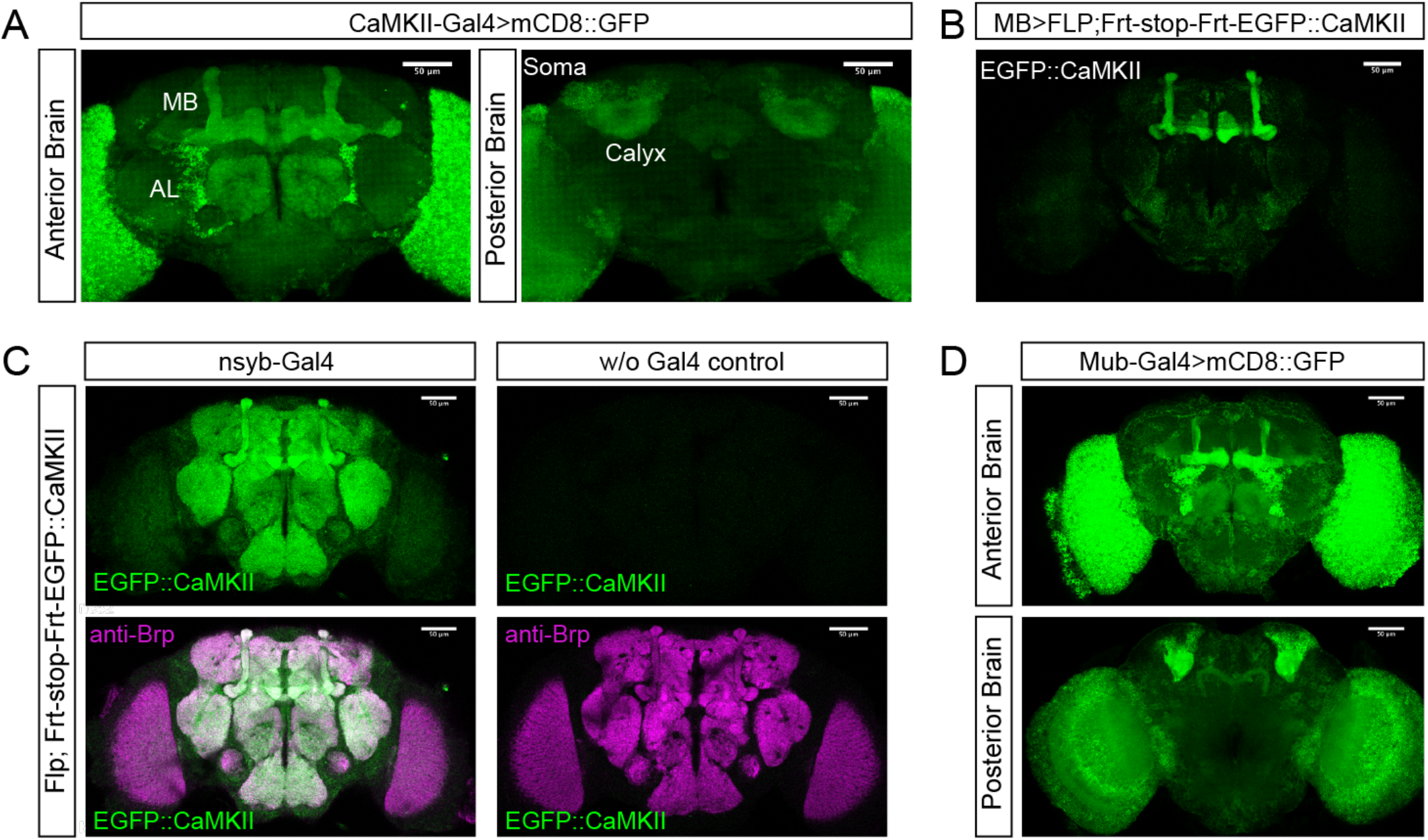
*CaMKII* and *Mub* genetic reagents, related to Figures 1 and 3. (A) The MiMIC transgene *Mi{MIC} CaMKII*^*MI03976*^, which is located in the ninth intron of the *CaMKII* gene was converted to a GAL4 line using standard techniques [33]. Confocal stacks of the anterior brain (left) and the posterior brain (right) from *CaMKII>mCD8::GFP* animals expressing membrane-bound GFP. (B) EGFP::CaMKII protein expression exclusively in Kenyon cells using *VT030559-GAL4* (denoted in this paper as “MB-GAL4”) to drive the FLP recombinase to remove stop sequences. (C) EGFP::CaMKII expression using pan-neuronal GAL4 to drive FLP. Left; *nsyb-GAL4* pan-neuronal flip out. Right; no EGFP signal is seen when no GAL4 is present. (D) The MiMIC transgene *Mi{MIC}mub*^*MI08161*^, which is located in the first intron of the *mub* gene, was converted to a GAL4 line using standard techniques. Confocal stacks of the anterior brain (top) and the posterior brain (bottom) from *mub>mCD8::GFP* animals expressing membrane-bound GFP show cells which transcribe *mub*. Scale bars = 50 µm for each panel.

**Figure S2.**
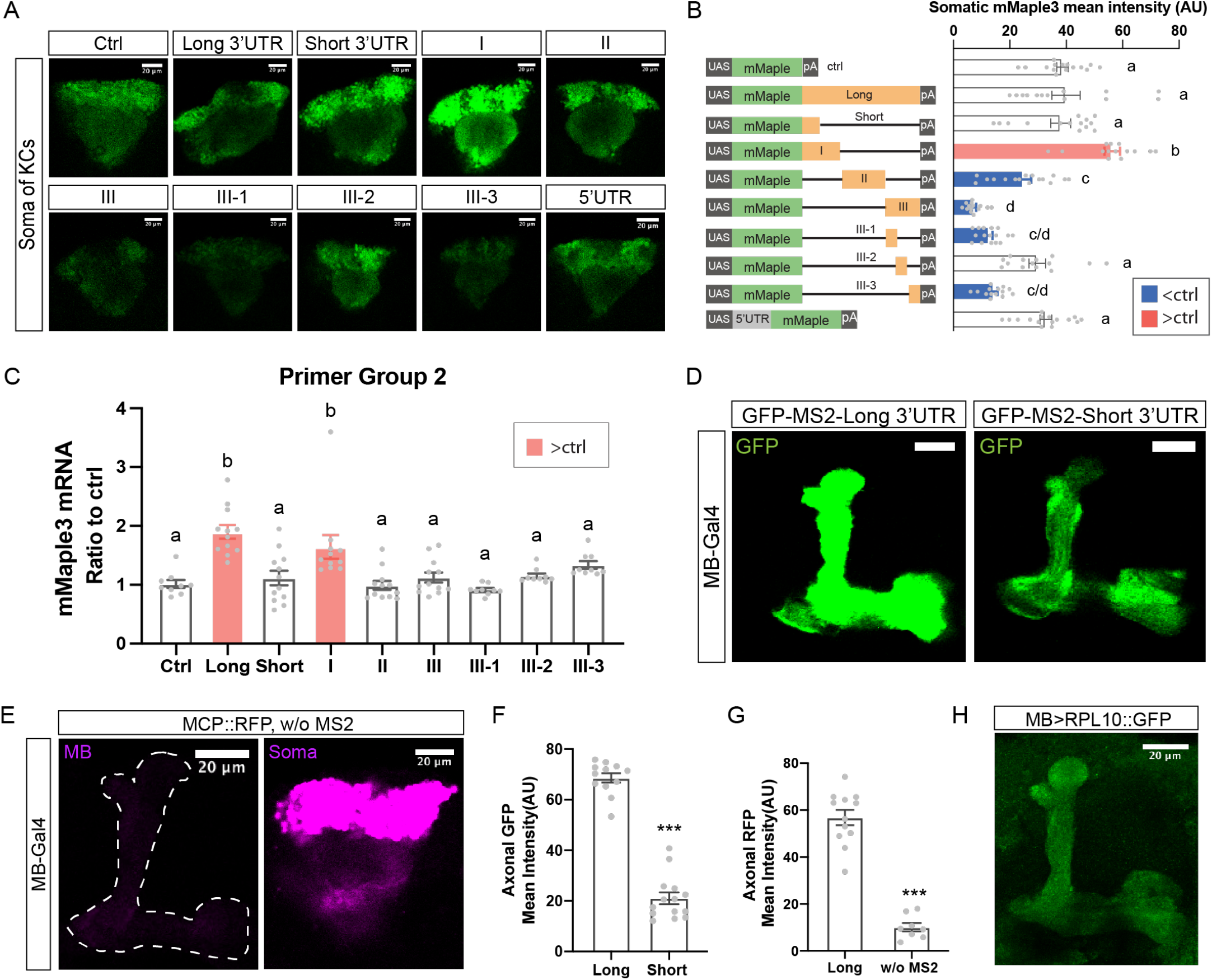
*Cis*-elements in the *CaMKII* 3’UTR which support axonal protein synthesis in MB, related to Figure 2. (A) Representative images of MB somatic mMaple3 protein expression from *MB>mMaple3* brains. Scale bars = 20 µm. (B) Left, cartoon of transgenes. Right, quantification of somatic mMaple levels and comparison to control transgene. N = 14-18. (C) qPCR of mRNA from adult *MB>mMaple3* heads using a second, independent primer set against mMaple3. N = 9-12. These data confirm that the proximal 3’UTR-I contains an mRNA stability element. For both panels B and C, red bars indicate significant increase relative to control, blue indicates decrease. (D) Representative images of the MB axonal region from *MB>24XMS2-myrGFP-UTR* animals. Left panel shows myrGFP protein reporter with long 3’UTR and right panel shows short 3’UTR. (E) Axonal MCP::RFP accumulation requires the expression of the MS2 transgene. Left panel shows MB axonal region from *MB>RFP::MCP-nls* animals that are not expressing an MS2-containg transgene. Right panel shows that all of the RFP::MCP-nls protein is sequestered in nuclei. (F) Quantification of myrGFP protein levels from panel D. N = 12-14. (G) Axonal RFP levels in animals with no MS2 transgene compared to animals co-expressing the *UAS-24XMS2-myrGFP* transgene. N = 8-12. (H) Ribosomes are present in the MB axonal compartment. Image shows MB region from *MB>RPL10::GFP* adult brain. This GFP-tagged ribosomal protein has been shown to co-assemble with endogenous ribosomes [15]. Data are shown as mean ± SEM and quantified by one-way ANOVA with Bonferroni post-hoc test or Student’s t-test accordingly. Statistical differences are indicated by letters in panels B and C; groups that are not significantly different have the same letter. *** indicates that P < 0.001 in panels F and G. Gray dots show individual values. Scale bars = 20 µm for each panel.

**Figure S3.**
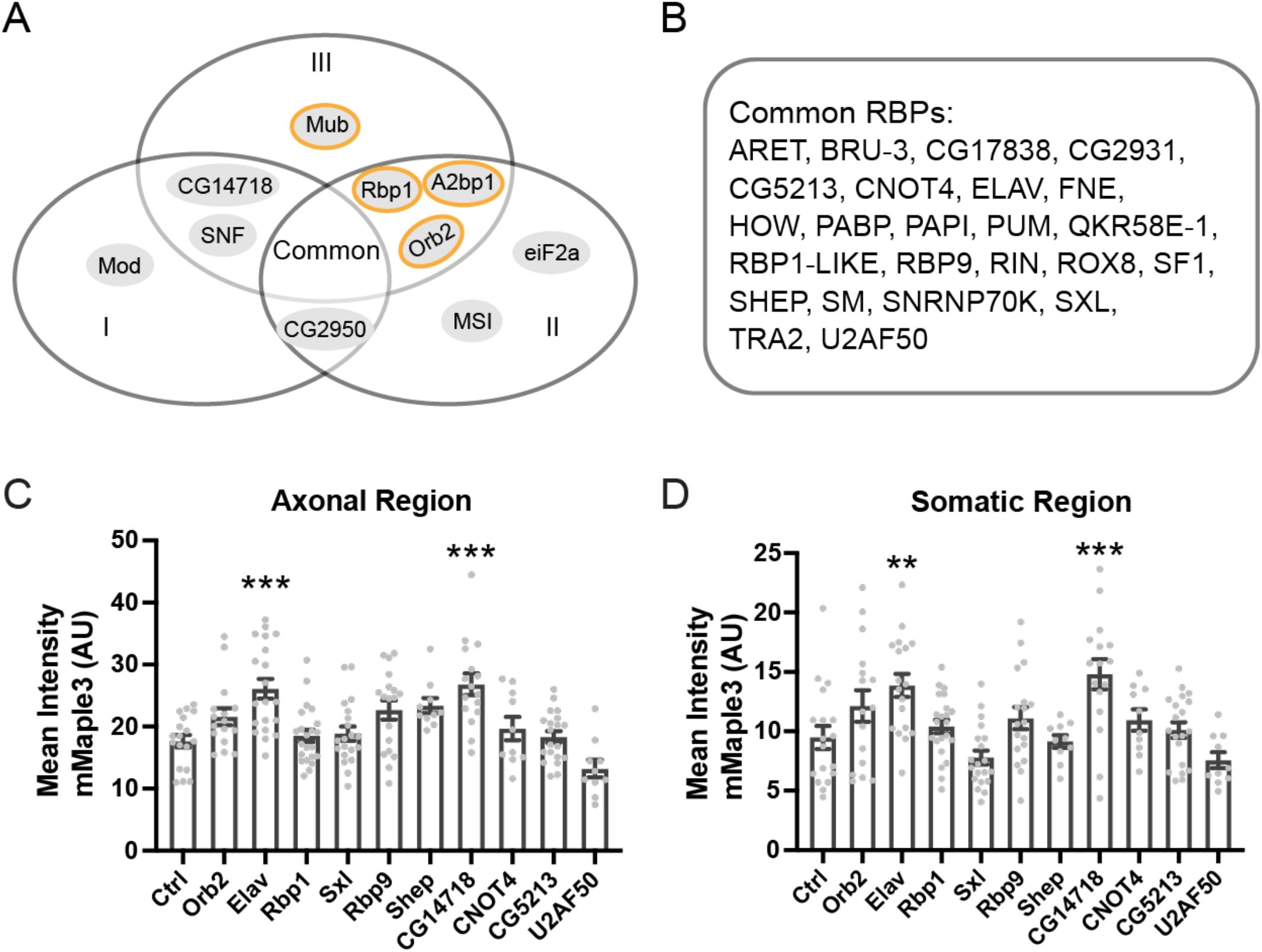
*In silico* predictions of RNA-binding protein and candidate *trans-*acting repressors, related to Figure 3. (A) The 3’UTR of the *CaMKII* gene was run through RBPmap [17] at medium stringency to obtain a list of potential RBP binding sites for screening. (B) Common RBPs found in all three fragments. (C/D) Candidate RBPs for screening were selected from the RBPmap list using the criteria that they must have at least one site in Fragment III and be predicted with P < E-04. RNAis for candidate RBP regulators were co-expressed in MB with *UAS-mMaple3-III*. Quantification of mMaple3 protein in *MB>mMaple3-III* ± RNAi brains shows that knocking down *elav* or *CG14718* significantly increased mMaple3 levels in both somatic and MB axonal regions. Data are shown as mean ± SEM and quantified by one-way ANOVA with Bonferroni post-hoc test. N = 10-22 for both panels. ** P < 0.01; *** P < 0.001. Gray dots show individual values.

**Figure S4.**
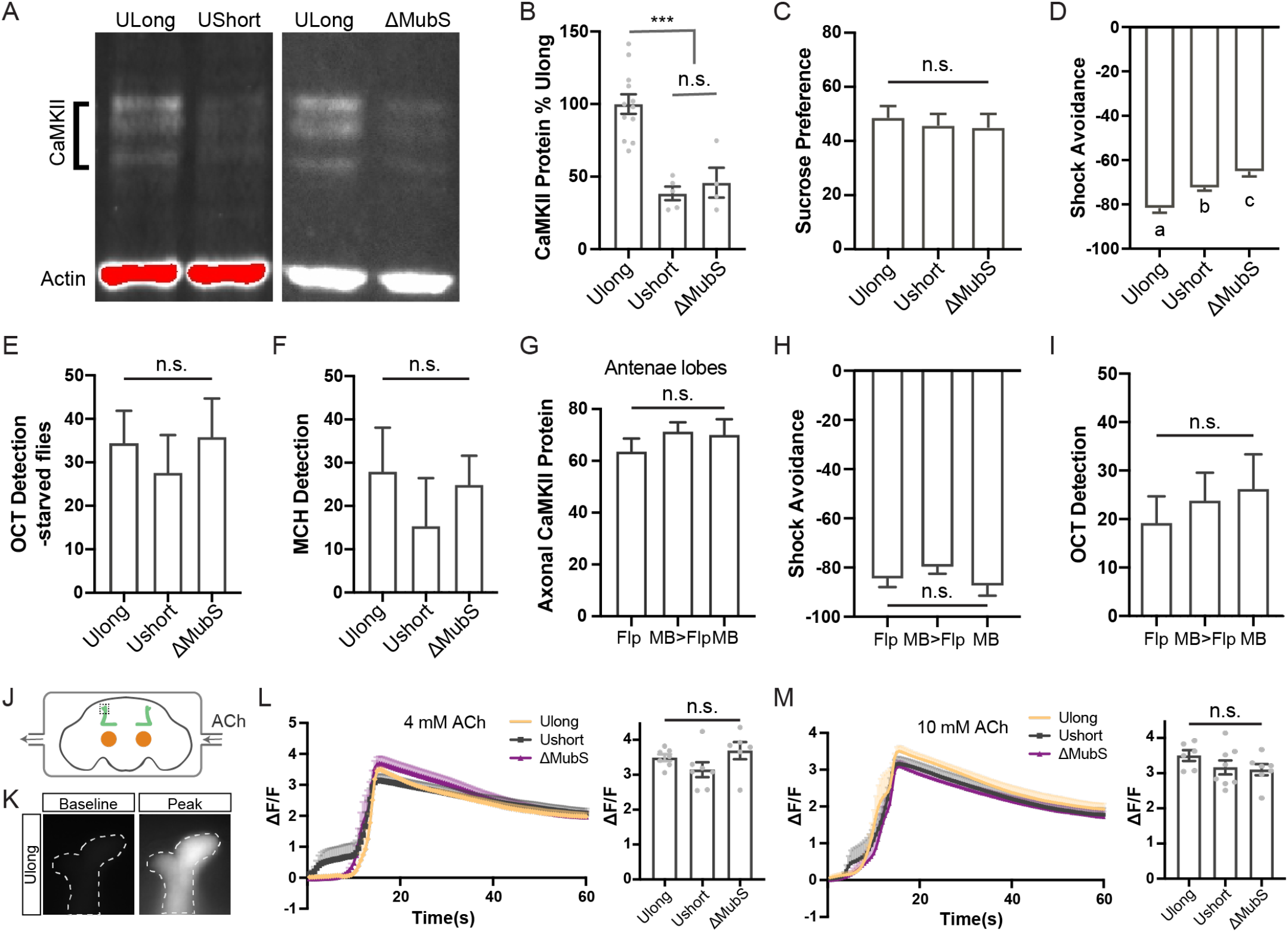
Loss of *CaMKII* 3’UTR reduces CaMKII protein but leaves odor and sucrose response unaffected, related to Figure 4. (A) Representative immunoblots with anti-CaMKII and anti-actin for normalization. SDS-PAGE for immunoblotting with monoclonal anti-CaMKII antibody showed that both *CaMKII*^*UShort*^ and *CaMKII*^Δ*MubS*^ have reduced total-head CaMKII compared to *CaMKII*^*ULong*^ which has the WT full length 3’UTR. (B) Quantification of CaMKII protein normalized to actin. N = 4-12. (C) *CaMKII*^*UShort*^ and *CaMKII*^Δ*MubS*^ show similar sucrose preference to *CaMKII*^*ULong*^. N = 6-9. (D) *CaMKII*^*UShort*^ and *CaMKII*^Δ*MubS*^ show similar shock avoidance to *CaMKII*^*ULong*^. N = 8 for each group. (E) *CaMKII*^*UShort*^, *CaMKII*^Δ*MubS*^ and *CaMKII*^*ULong*^ show similar ability to detect octanol, even when starved N =10-16. (F) *CaMKII*^*UShort*^, *CaMKII*^Δ*MubS*^ and *CaMKII*^*ULong*^ show similar ability to detect MCH, N = 8. (G) Quantification of CaMKII in antennae lobes showed that CaMKII protein level was unaffected when 3’UTR was specifically knocked out in MB. N = 16 for each group. (H) MB-specific knockout of *CaMKII* 3’UTR sequences didn’t affect shock avoidance. N = 8-12. (I) MB-specific knockout of *CaMKII* 3’UTR sequences didn’t affect octanol detection. N = 10-26. (J) Diagram of ACh perfusion setup. Different concentrations ACh were perfused across the imaging region. The dashed square shows the recording area, the α/α’ tip region. (K) Representative pictures of the recording area for baseline and peak calcium responses of *ULong* flies (*MB/20xGCaMP6f*; *CaMKII*^*ULong*^). (L) With 4 mM ACh perfusion, *CaMKII*^*UShort*^ and *CaMKII*^Δ*MubS*^ flies showed comparable calcium responses to *CaMKII*^*ULong*^. N = 6-7. (M) With 10 mM ACh perfusion, calcium responses of *CaMKII*^*UShort*^ and *CaMKII*^Δ*MubS*^ were also similar to that of *CaMKII*^*ULong*^. N = 6-8. The genotypes in panels L and M are: *MB/20xGCaMP6f*; *CaMKII*^*ULong*^, *MB/20xGCaMP6f*; *CaMKII*^*UShort*^ and *MB/20xGCaMP6f*; *CaMKII*^Δ*MubS*^. Data are shown as mean ± SEM and quantified by one-way ANOVA with Bonferroni post-hoc test. *** P < 0.001, and n.s. indicates no significant difference. Statistical differences in panel D are indicated by letters; groups that are significantly different have different letter.

**Table S1.**
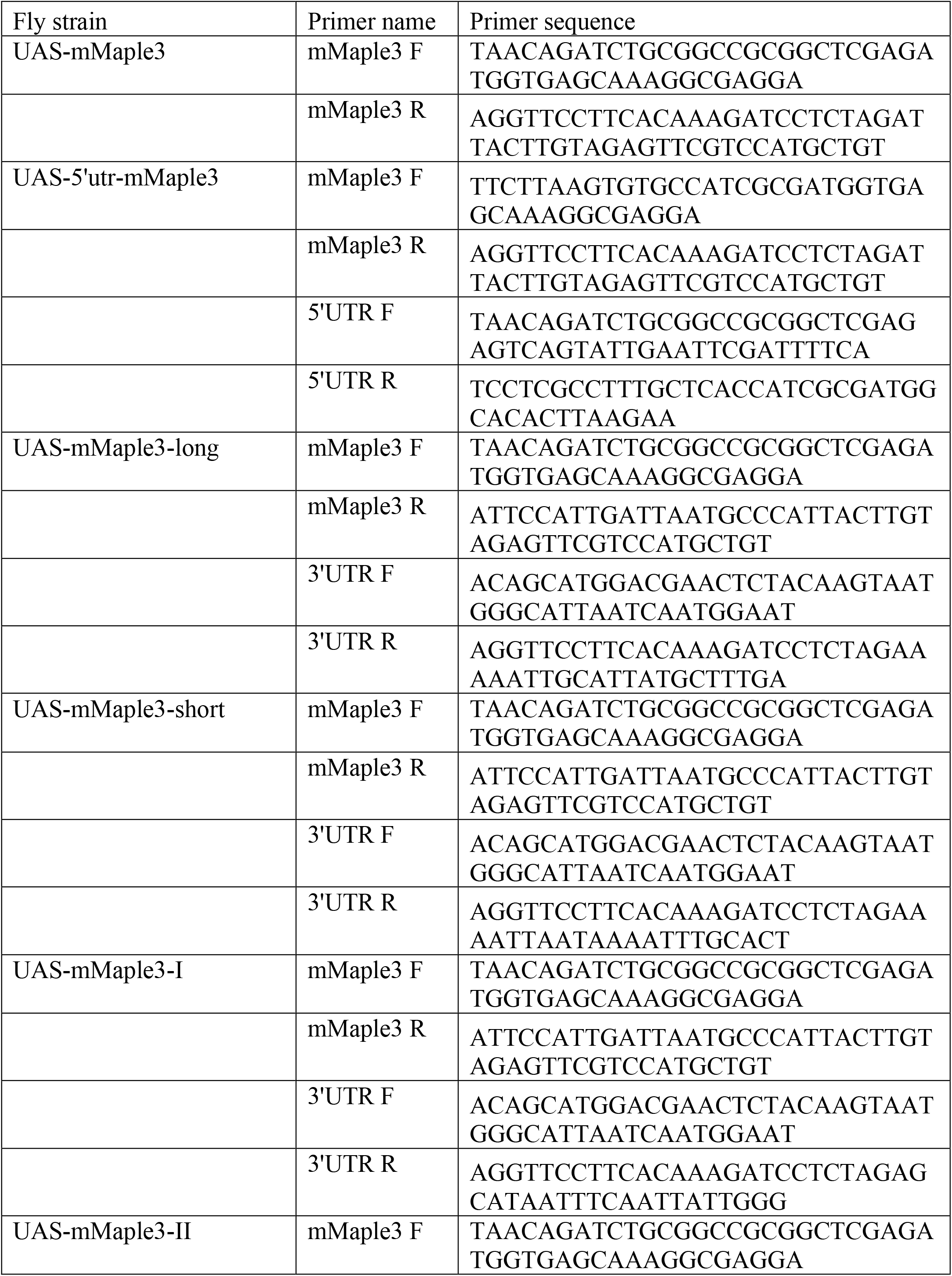
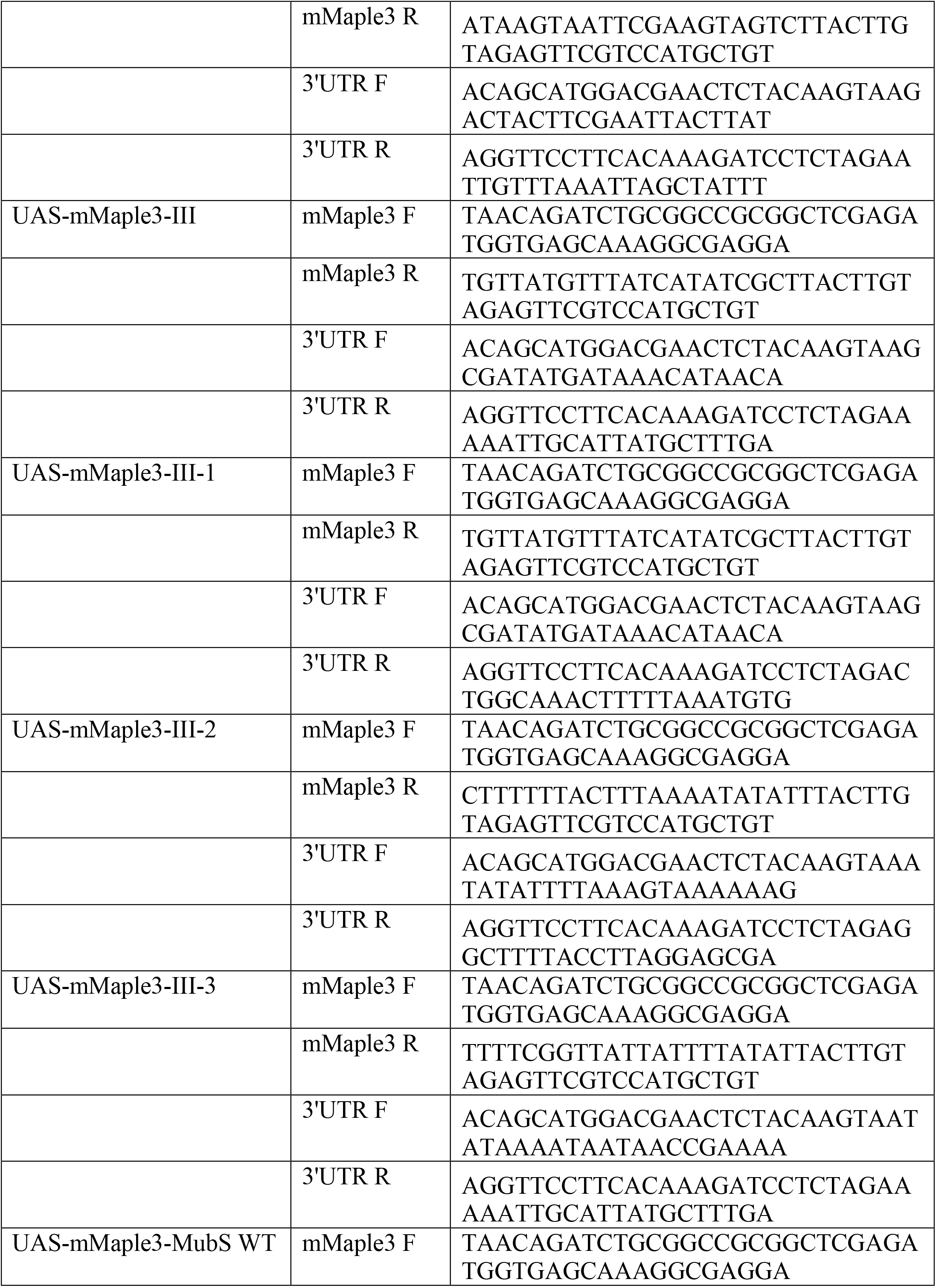
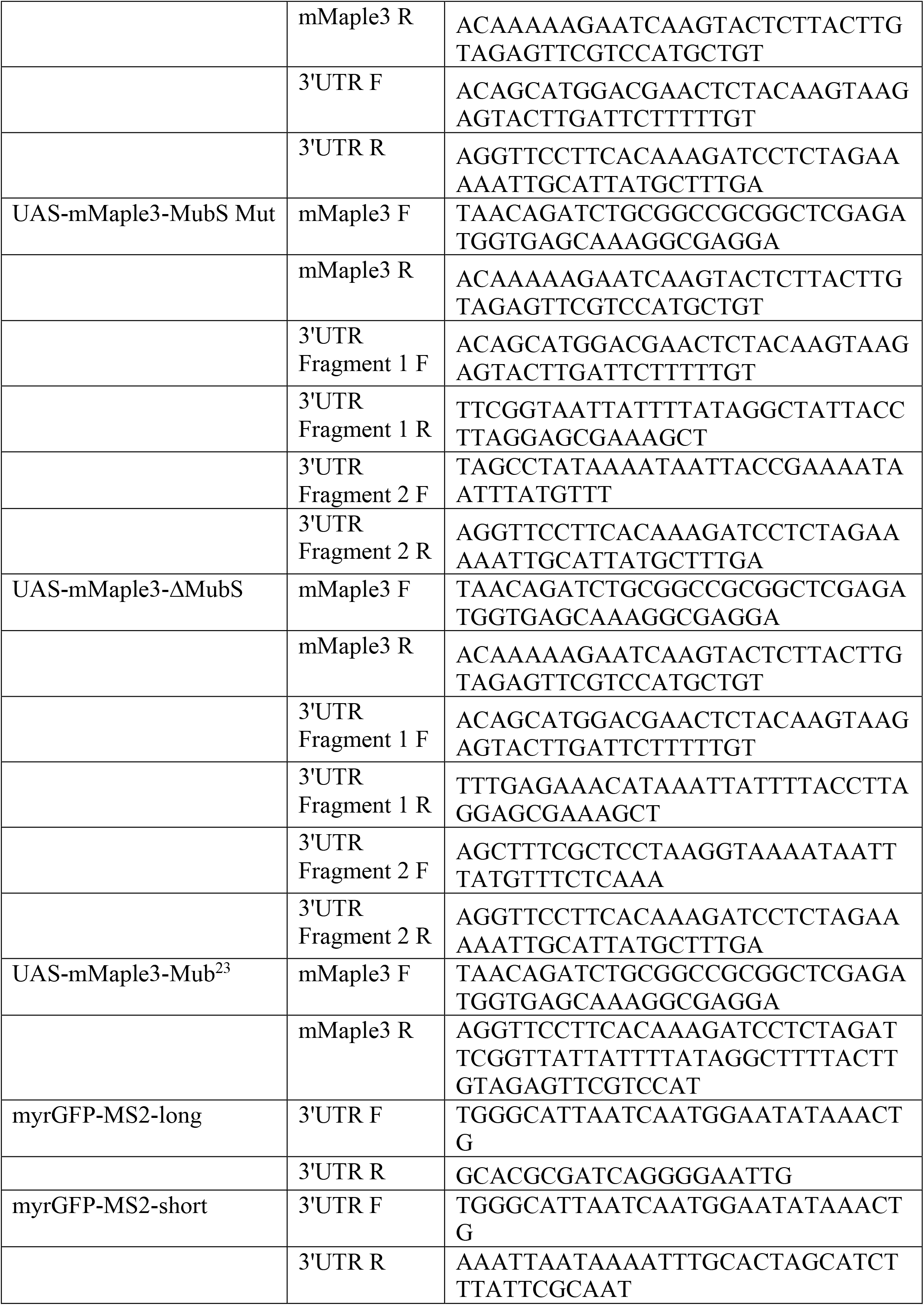
Primer list for transgenic flies.

**Table S2.**
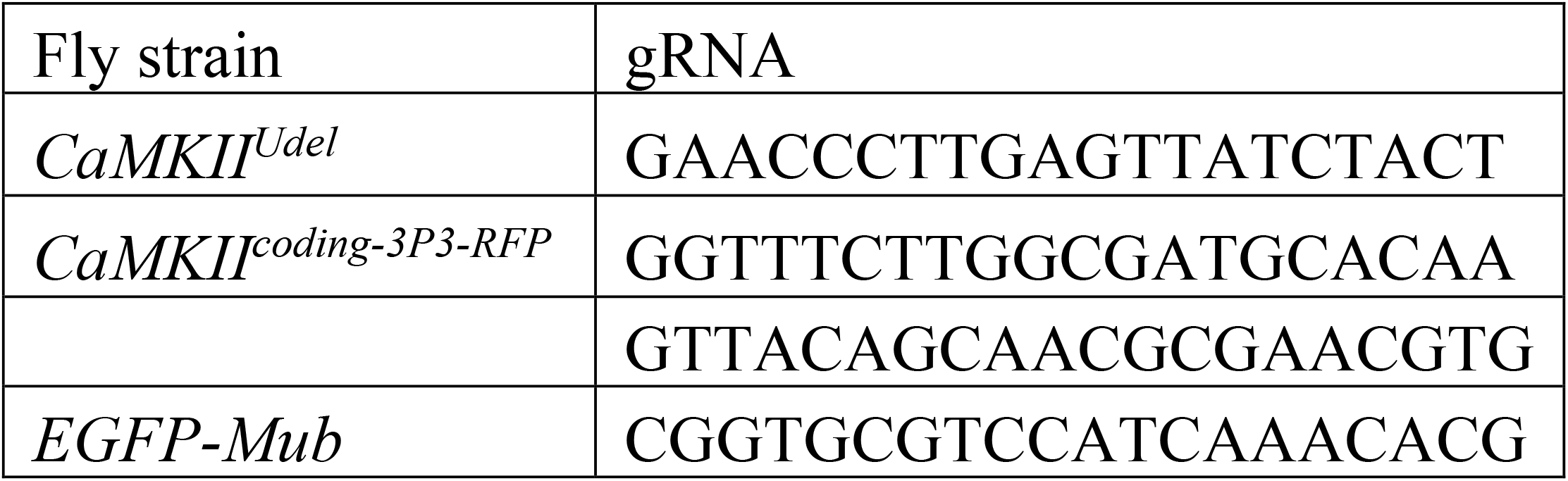
Guide RNA list for CRISPR/CAS9 flies.

**Table S3.**
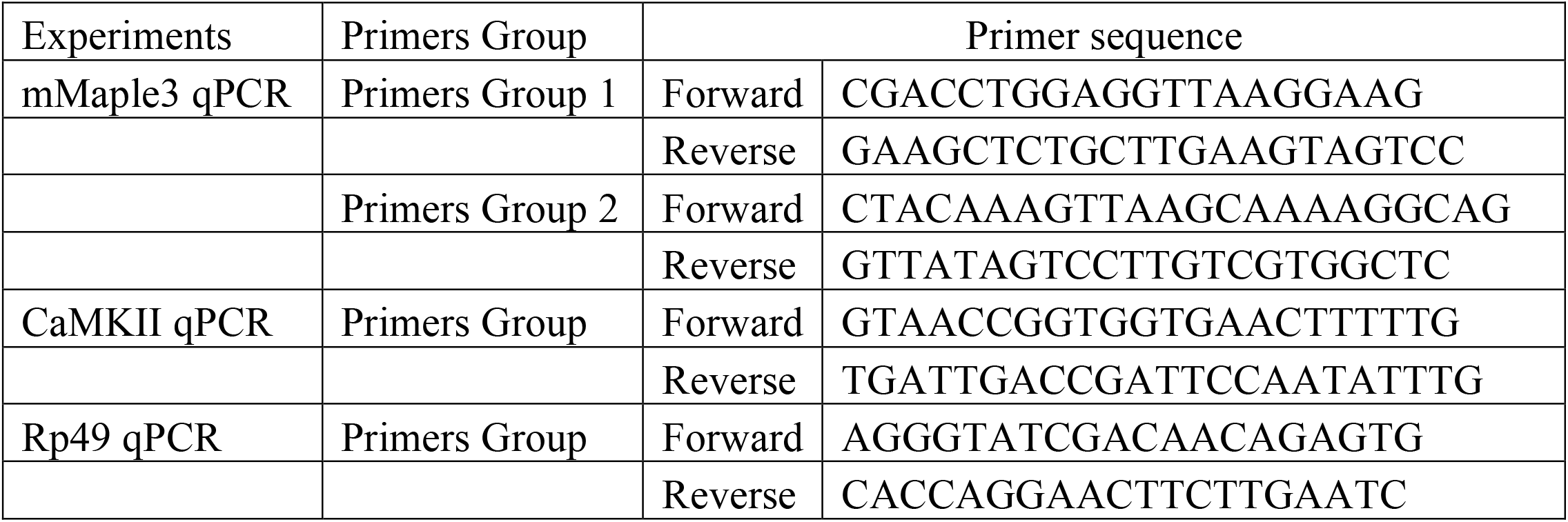
Primer list for qPCR experiments.

## Notes

### Competing Interest Statement

The authors have declared no competing interest.

